# Controllable membrane damage by tunable peptide aggregation with albumin

**DOI:** 10.1101/2022.06.02.494564

**Authors:** Seren Hamsici, Gokhan Gunay, Handan Acar

## Abstract

Aggregation of otherwise soluble proteins into amyloid structures is a hallmark of many disorders, such as Alzheimer’s and Parkinson’s diseases. There is increasing evidence and acceptance that instead of ordered amyloid assemblies, the misfolded oligomer aggregations are the main toxic structures. However, there is no system to study the mechanism and kinetics of aggregation, distinguish between ordered structures and misfolded oligomers, and correlate the structures to their toxicity. The exact role of oligomer aggregation in the pathological process remains to be elucidated. Here, we use an engineered co-assembling oppositely charged amyloid-like peptide pair ([II]) to relate its aggregation to toxicity. The toxicity mechanism of [II] is through cell membrane damage and stress, as shown with YAP and eIF2α biomarkers, as in the amyloid protein-initiated diseases. Albumin is used to control the aggregation of [II], and so its toxicity. This study represents a molecular engineering strategy to study the aggregation process of amyloid-like structures in diseases. Understanding the nature of protein aggregation through engineered peptides paves the way for future designs and drug development applications.

## 1. INTRODUCTION

A wide range of proteins aggregate into amyloids, cross β-sheet fibrillar structures that are linked to numerous diseases, such as Alzheimer’s and Parkinson’s. Although for many decades the accumulation of amyloid plaques was associated with the pathology of the diseases, increasing evidence suggests that the misfolded protein aggregates, oligomers, that are soluble and form in the process of fibrillization, are the main cause of the toxicity.^1–4^ Part of their toxicity process occurs through the interactions of oligomers of amyloid-β or α-synuclein with the lipid membranes of cells, which initiates damage.^5,6^ The membrane damage often induces immunogenic cell death and inflammation.^7–9^ The aggregation and oligomer formation process is complex because of the convoluted conditions in the body. The dynamic conditions create transient heterogeneous misfolding of oligomers and make the molecular level of understanding, quantifying, and correlating the aggregation mechanism and toxicity, and thus developing drug targets, challenging.^4,10^ Therefore, a minimalist and simplified synthetic model that mimics the structure, aggregation, and function of amyloids can serve as a powerful tool for studying their aggregation mechanism.

Co-assembly of oppositely charged peptides (CoOP) is a minimalist strategy enabling to study the role of intermolecular interactions in peptide aggregation.^11^ CoOPs are designed to assemble into amyloid fibrils. Among the studied peptide sequences, the oppositely charged pair, KFFIIK and EFFIIE (the pair will be shown henceforth as [II]), showed the most ordered amyloid-like β-sheet assembly. On the cell membrane, [II] peptide-aggregation induces immunogenic rupture (PAIIR) that initiates immunogenic cell death (ICD) and associated release of damage associated molecular patterns (DAMPs) as an immunostimulatory tool.^12^ PAIIR induces secondary pyroptosis, a programmed ICD and a common feature of amyloid inducing cell death as in Alzheimer’s and Parkinson’s diseases.^13,14^ PAIIR is a simplified peptide-based tool, designed to mimic the function, instead of the sequence of a protein; thus there is no natural ortholog of [II].

Albumin is abundant in the human body, flexible, and mostly hydrophobic, and thus, it has high non-specific binding capacity.^15^ It is known that albumin impedes the fibrillization of amyloids and increases the soluble misfolded oligomers of higher toxicity.^16–18^ Akin to amyloid, co-aggregation with albumin creates a soluble oligomeric form of [II] that is more toxic than its fibril form. In this study, we show control over the aggregation kinetics and mechanism of [II] via albumin. The simplicity of [II] outlines the detailed characterization of aggregates and their interactions with the lipid membranes. The lipid membrane interaction is shown with liposomes and fluorescent tagged peptides. The soluble form of peptides misfolded with albumin integrate into the lipid membrane of liposomes, while the ordered fibril structures do not. The kinetics of the aggregations are studied with spectroscopic methods with fluorescent probes. The morphology of misfolded oligomeric to ordered fibrillar structures are analyzed by dynamic light scattering (DLS) and atomic force microscopy (AFM). In parallel with membrane localization, cellular cytotoxicity follows a similar trend; the oligomeric form of [II] misfolded with albumin causes membrane damage and cellular cytotoxicity, while the fibrillar [II] without albumin does not. By delaying the formation of thermodynamically favorable ordered fibrils via albumin or simply introducing the counterparts in different time points, the soluble free peptides are able to localize into the membrane before they come together. Lastly, the mechanical cell stress induced by [II] is identified through the phosphorylation of yes-associated protein (YAP) and eukaryotic initiation factor 2 alpha (eIF2α), similar to the mechanism of amyloid-β cytotoxicity. Importantly, we identified these stress markers in the cells treated with misfolded oligomers, not with ordered amyloid fibrillar form of [II]. Overall, [II] presents a simplistic tool to study the effects of physical and biological processes to understand aggregation process and function of amyloid-like pathological proteins.

## 2. Results and Discussion

### 2.1 Integration of [II] peptides into liposomal membrane

Liposomes are commonly used tools for the analysis of interactions between amyloids and lipid membranes.^19,20^ We used the agarose embedded liposomes (blue, app. 100 μm in diameter, **Figure S2**) for long time imaging and the fluorescence tagged [II] peptides (KFFIIK with FITC (green) and EFFIIE with rhodamine B (red)) to identify their membrane interactions **(Figure 1A)**.^21^ 30 min premixed [II] as in **Figure 1A-i** initiates the interaction of oppositely charged groups, as the charge neutralization was observed through zeta potential **(Figure S3)**. The high affinity of peptides prevented disassembly and localizing on the lipid membrane of liposomes **(Figure 1B)**.

**Figure 1.**
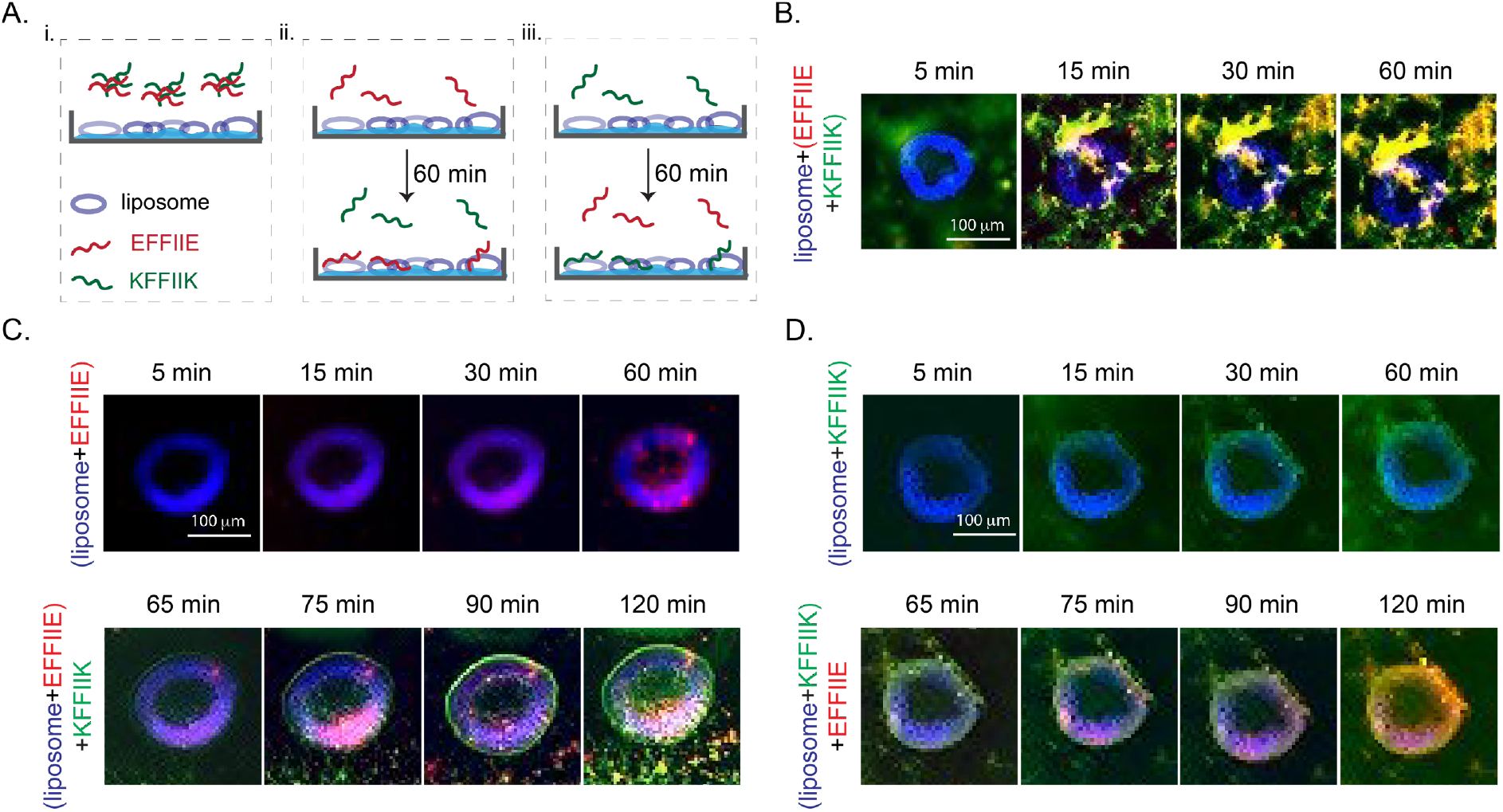
Liposome (blue) membrane integration of [II] and individual peptides; KFFIIK (green), EFFIIE (red). (A) The schematic of membrane localization experiment by using agarose embedded liposomes. (B) Membrane localization of 30 min premixed of EFFIIE+KFFIIK. (C) EFFIIE addition followed by KFFIIK and (D) KFFIIK addition followed by EFFIIE. (Scale bar = 100 μm).

Incubation of EFFIIE alone for 60 min as in **Figure 1A-ii** allowed the integration of the peptide into the cell membrane (red fluorescence on blue liposomes). Despite the charge similarities with the lipid, the hydrophobicity of the FFII group might be the reason of this integration. After introducing KFFIIK on the EFFIIE+liposome, green fluorescent (KFFIIK) increased immediately in 5 min, which shows the localized interaction of [II] on the membrane **(Figure 1C)**. Additionally, amyloid fibril aggregation around the liposomes was also observed, indicating that the remaining soluble EFFIIE interacts with its soluble counterpart peptide before integration into the lipids. Similarly, incubation of KFFIIK alone, as in **Figure 1A-iii**, showed integration into the lipids **(Figure 1D)**. The membrane localization of KFFIIK was faster than EFFIIE, likely because of the opposite liposome surface charge.^22^ Addition of EFFIIE 60 min later also showed membrane integration of both peptides and also fibrillar aggregations of [II] outside of the membrane.

Overall, these results demonstrated that the soluble forms of individual [II] peptides integrate into membrane. The premixed [II] did not show peptide integration; they were stable. The membrane-binding capacity of [II] is correlated to the presence of free peptides in the medium. Controlling the aggregation kinetics of [II]—in another words, controlling the time and amount of free peptides in the medium—can control the amount of peptides integrated into the membrane.

### 2.2 Controlling peptide aggregation kinetics

The mechanism of ordered fibrillar amyloid assembly follows sigmoidal growth divided into three steps: (i) lag phase (nucleation) in which the peptides are mostly in misfolded monomeric form; (ii) growth (elongation) phase in which the larger aggregates (misfolded or ordered) form of peptides and (iii) final plateau in which grown aggregates (misfolded or ordered) dominate at the final equilibrium.^23^ Amyloid forming proteins have hydrophobic domains which can misfold and form non-specific interactions and disordered aggregates with the other hydrophobic molecules in the body. The stability of these aggregates depends on the affinity of each component. Because ordered amyloid assembly of peptides have specific interactions that require a specific conformation, they are often more stable than non-specific hydrophobicity driven misfolded aggregates. However, the same conformational specificity creates an energy barrier. Any aggregation in the intermediate energy level, particularly when it is highly non-specific, impedes the conformational order for the assembly and delays the equilibrium in the final plateau of stable fibrils. In other words, misfolding of monomers prolongates the initial lag and growth phase where the monomeric structures are soluble and less stable.

Albumin, as a flexible hydrophobic protein, aggregates with amyloid forming proteins, prolonging the lag phase.^16–18^ The unstable and soluble monomers present in the lag phase are able to integrate to the cell membrane and become more toxic.^24^ Although the prolonged lag time is known to be more toxic, there has been little study on the correlation between the lag time (aggregation kinetics), structures of the aggregates, and cell membrane damage. Since [II] showed amyloid-β characteristics such as highly ordered secondary structure, we hypothesized that albumin can change the aggregation behavior and fibrillar assembly kinetics of [II].^11^ **Figure 2** shows the kinetics and morphology of [II] aggregates in increasing amount of albumin (0, 0.002, 0.02, 0.05, 0.1, 0.2, 0.5, 1, and 2 w%) in PBS. To distinguish the kinetics of ordered amyloid assembly from misfolded oligomer aggregates, we used Thioflavin T (ThT) and 1,6-diphenyl-1,3,5-hexatriene (DPH) assays, respectively. The morphology and size of the aggregates were studied with AFM and DLS.

**Figure 2.**
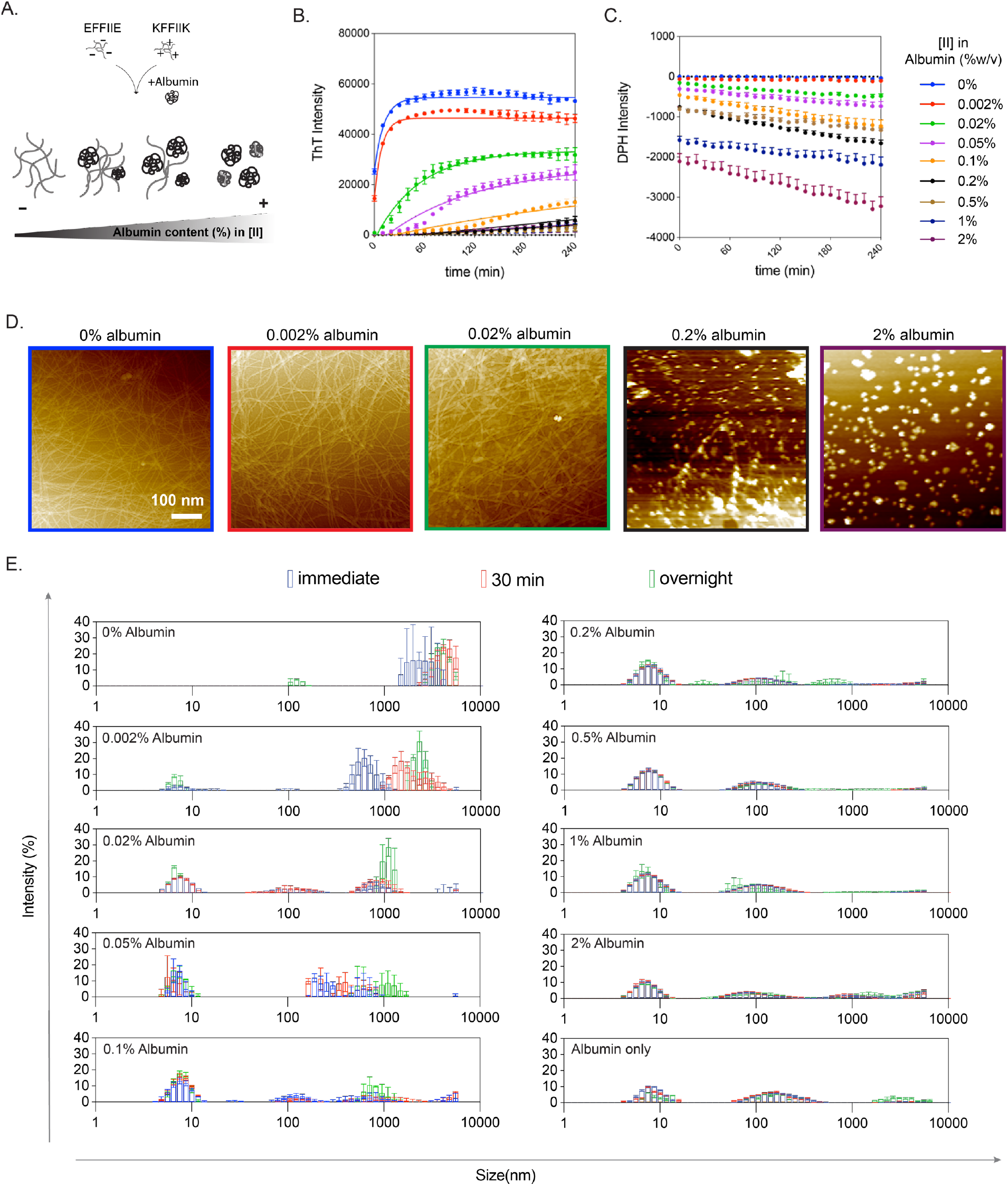
Aggregation kinetics depend on albumin concentration. (A) The schematic of [II] aggregation with albumin. Aggregation kinetics were performed with (B) ThT and (C) DPH assays in increasing albumin content. (D) AFM images of [II]-albumin structures (Scale bar = 100 nm). (E) DLS measurements [II]-albumin mixtures.

ThT is a fluorescent dye commonly used for β-sheet structure identification.^25^ The transition from oligomers to fibrils has been characterized with ThT in prion protein;^26^ here, we apply it to [II] Individual peptides were prepared in PBS with increasing albumin content and constant ThT, and the fluorescence of ThT was tracked for 4h **(Figure 2B)**. Albumin only groups with different w% were measured with constant ThT as the background, which also showed some degree of intensity as the structures have hydrophobic core, but did not increase with time **(Figure S4)**. When mixed, [II] peptides assemble into ordered structures and achieve their final equilibrium of fibrillar form in less than 10 min.^11^ Fibrillar final equilibrium was only observed in the medium without albumin (0 w%) and 0.002, 0.02, 0.05, and 0.1 w% albumin in 4h **(Figure 2B)**. Similarly, the fibrillar structures were observed in the overnight incubated samples in these parameters with AFM **(Figure 2D)**. The addition of albumin delayed the formation of ordered amyloid structures by increasing the rate of the lag phase (nucleation) and growth (elongation) phase ^27^. There was also a reduction in the total amount of fibers generated as indicated by ThT fluorescence signals as shown with AFM, and no fibrillar structure was observed for 0.2 and 2 w% albumin incubated overnight **(Figure 2D)**.

The formation of disordered structures was identified with DPH assay; DPH is a fluorescent dye with enhanced intensity in hydrophobic environments.^28^ Because albumin itself is a hydrophobic protein and forms soluble aggregates in PBS, the initial measurements were performed with only DPH and albumin as the background **(Figure S5)**. Each peptide was prepared in PBS with concentrations increasing in albumin and constant in DPH, then mixed and tracked for 4h **(Figure 2C)**. The decreased intensity of DPH with increasing albumin is interpreted as the decrease the number and sizes of the aggregates. This might be due to the covering of the hydrophobic parts of the albumin with the charged peptides creating more soluble and smaller albumin aggregates in PBS. In the PBS with the lowest amount of albumin, 0.002 w%, albumin showed an insignificant intensity and the structures in this solution are mostly fibrillar, as observed through increasing ThT assay, indicating that the amount of disorganized aggregates were negligible. Presence of increased albumin concentrations showed slowly lowered intensities; in other words, more change between the intensity of the [II] + albumin complexes compared to the same amount of albumin alone, is because of formation of smaller aggregates.

While the aggregation of individual peptides did not change with the addition of albumin, several observations are worth noting. When alone, KFFIIK showed slightly higher and more stable ThT intensity, which reduced but remained stable over time with increasing albumin **(Figure S4)**. Similarly, the DPH assay showed increasing intensity over time in KFFIIK alone **(Figure S6)**. These measurements were all performed in PBS in which the charges of the peptides are screened by the salt ions. Therefore, KFFIIK alone shows misfolded aggregation due to the hydrophobic nature of the alkyl tail of Lysine (K). Yet, when KFFIIK is mixed with EFFIIE, the ThT intensity changes immediately and dramatically, indicating that the peptides co-assemble in PBS **(Figure 2B)**. The negative counterpart, EFFIE, in PBS without albumin, does not aggregate in an ordered or disordered manner **(Figure S4 and S6)**. Considering albumin alone also has ThT and DPH intensities due to its hydrophobic nature **(Figure S4 and S6)**, the observed consistent negative intensities of all EFFIIE and KFFIIK solutions with albumin indicates formation of smaller and more stable aggregates; peptides interact and aggregate with albumin individually. However, when mixed, the increase of ThT intensities, or formation of peptide fibers, indicates that the individual peptide and albumin aggregations are not as favorable as peptide to peptide interactions. Addition of albumin to premixed [II] showed stable DPH intensity lower than that of albumin only (**Figure S5**, no background subtraction), possibly because albumin falls apart and forms non-specific bindings with the fibrillar structures.

Size measurements of the aggregates was performed with DLS in three different time points: immediately, after 30 min, and after overnight incubation **(Figure 2E)**. Without albumin, [II] aggregated immediately with >1000nm average size with a shift toward higher sizes after 30 min (**Figure 2E**, 0 w% albumin). Similarly, the shift toward bigger structures was observed for the conditions between 0.002 to 0.1 w% albumin, which are the conditions where the final fibrillar equilibrium was observed with ThT. After 0.2 w%, size distribution and intensity profiles were similar to the 2 w% albumin only group. The population of structures around 10 nm were higher in the presence of peptides compared to 2 w% albumin only, indicates smaller aggregates in the presence of peptides, and explains the reduction in intensities observed with DPH.

Overall, the observed trend of [II] with albumin is similar to that of natural amyloids; the increasing albumin causes the decrease of the maximum ThT fluorescence intensity, or the total amount of fibrils generated, and increases soluble disordered aggregates; oligomers.

### 2.3 Delayed aggregation increased the lipid membrane integration of [II]

Among the studied albumin concentrations, 0.2 w% showed the threshold for diminishing the ordered fibril formation; hence, the rest of the analyses were performed in 0.2 w% albumin. To compare the integration on the lipid membrane of the liposomes, [II] prepared in 0.2 w%, incubated for 30 min, then added on the liposomes **(Figure 3A)**. Unlike the premixed [II] in 0 w% albumin **(Figure 1B)**, the individual peptides were localized in the liposomes starting from 5 min. Albumin slows down the peptide-peptide interactions by aggregating with the membrane, and the interactions of peptides with the lipid membrane is likely to be more stable than their interactions with albumin. Incubation of individual peptides with 0.2 w% albumin for 60 min showed that KFFIIK integrates to the lipid bilayer even in 5 min, while the negative counterpart, EFFIIE, does not integrate, likely due to the charges of the lipids **(Figure S7)**. The integration of EFFIIE when incubated with albumin and KFFIIK might be because of initial membrane integration of soluble KFFIIK followed by the aggregation of EFFIIE, even in the lipid interface. These results indicate that without albumin, KFFIIK and EFFIIE co-assemble into fibers, which do not interfere with the integrity of the lipid membrane. In the presence of albumin, the co-assembly process is prolonged due to the aggregation of the individual peptides with albumin. The delay time (lag time or the time required to achieve equilibrium fibril phase) is correlated to the amount of albumin in the solvent.

**Figure 3.**
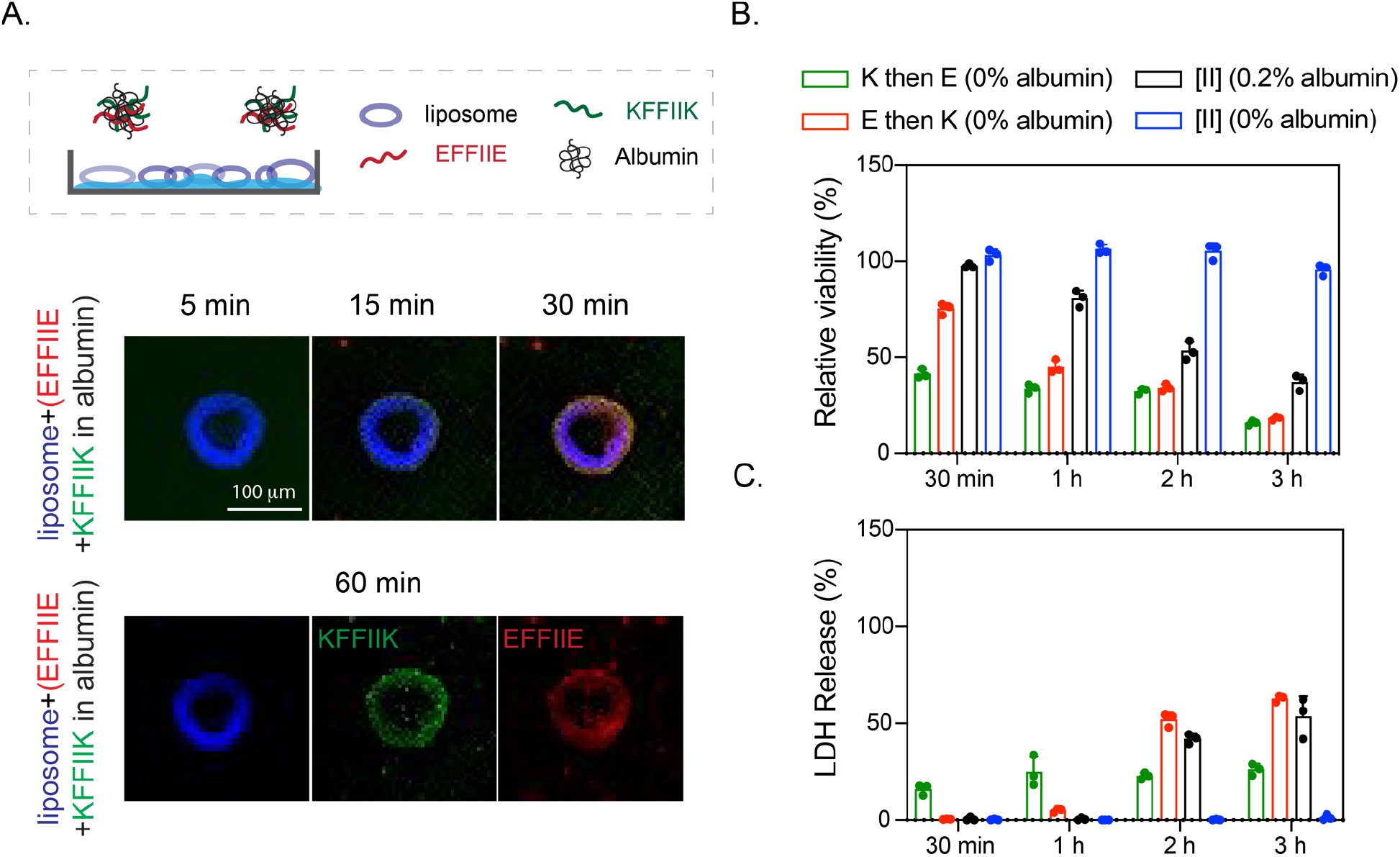
Delayed [II] aggregation increases its integration on the lipid membranes. (A) Liposomal membrane localization of 0.5 mM [II] in 0.2w% albumin; liposome = blue, EFFIIE = red, KFFIIK = green (Scale bar = 100 μm). (B) Relative viability and (C) LDH release of EMT6 cells treated with individual and aggregates with 0.5 mM [II] in 0 and 0.2 w% of albumin in PBS.

The effect of enhanced lipid membrane accumulation of [II] altered its toxicity **(Figure 3B, C)**. [II] induces immunogenic rupture of mammalian cell membranes, initiates immunogenic cell death in 6h in cell medium supplemented with 10 v% FBS (has app. 0.2w% albumin) as shown before.^12^ During 3h of treatment, [II] in 0 w% (without albumin) did not cause any membrane damage **(Figure 3B)**, as [II] fibrils do not interact with the lipid membrane **(Figure 1B)**. Addition of [II] with 0.2 w% albumin showed increasing membrane damage over time which is quantified by the release of lactate dehydrogenase (LDH), a cytoplasmic enzyme released upon membrane damage.^29^ These results correlate with ThT (high intensity, high ordered fibril assembly) and DPH results (low intensity, high disordered aggregation) indicating that delayed fibril formation increases the membrane damage. Furthermore, we also tested the effects of consecutively introduced peptides to cells without albumin to correlate the toxicity to the membrane integration. When KFFIIK was given first (green bar) followed by EFFIIE addition, cellular viability decreased to 50% at 30 min and reduced to 25% at 3h **(Figure 3B)**. The opposite scenario where EFFIIE added first and then KFFIIK (red bar), started from 75% cell viability at 30 min and decreased to 25% at 3h. As observed from liposomal experiments **(Figure 1C, D)**, KFFIIK accumulated faster on the membrane compared to EFFIIE. Therefore, when peptides were introduced consecutively (30 min), we observed faster cell death in K then E condition. However, cell death was similar at 3h regardless of the addition order **(Figure 3B)**. These results show that the toxicity of [II] can be controlled through control over its aggregation kinetics.

### 2.4 FRET analysis for albumin and [II] interactions

FITC and TRITC are fluorescent dyes and FRET couples. When the distance of the two fluorophores is less than 8 nm, the excitation of light with 488 nm wavelength triggers FRET; emission from FITC (donor) at 520 nm excites TRITC (acceptor) that emits light at 578 nm.^30^ We used albumin conjugated with TRITC and incubated with FITC-KFFIIK for this analysis. Incubation of FITC-KFFIIK with 0.2 w% albumin for 30 min and 24h which did not show FRET, indicates that they are not in close proximity **(Figure 4A)**. In the presence of EFFIIE (not fluorescent tagged) also in 0.2 w% albumin showed the FRET intensity, TRITC emission at 580 nm and FITC emission at 488 nm was reduced **(Figure 4B)**, indicating the peptides and albumin are aggregated in closer proximity than 8 nm.

**Figure 4.**
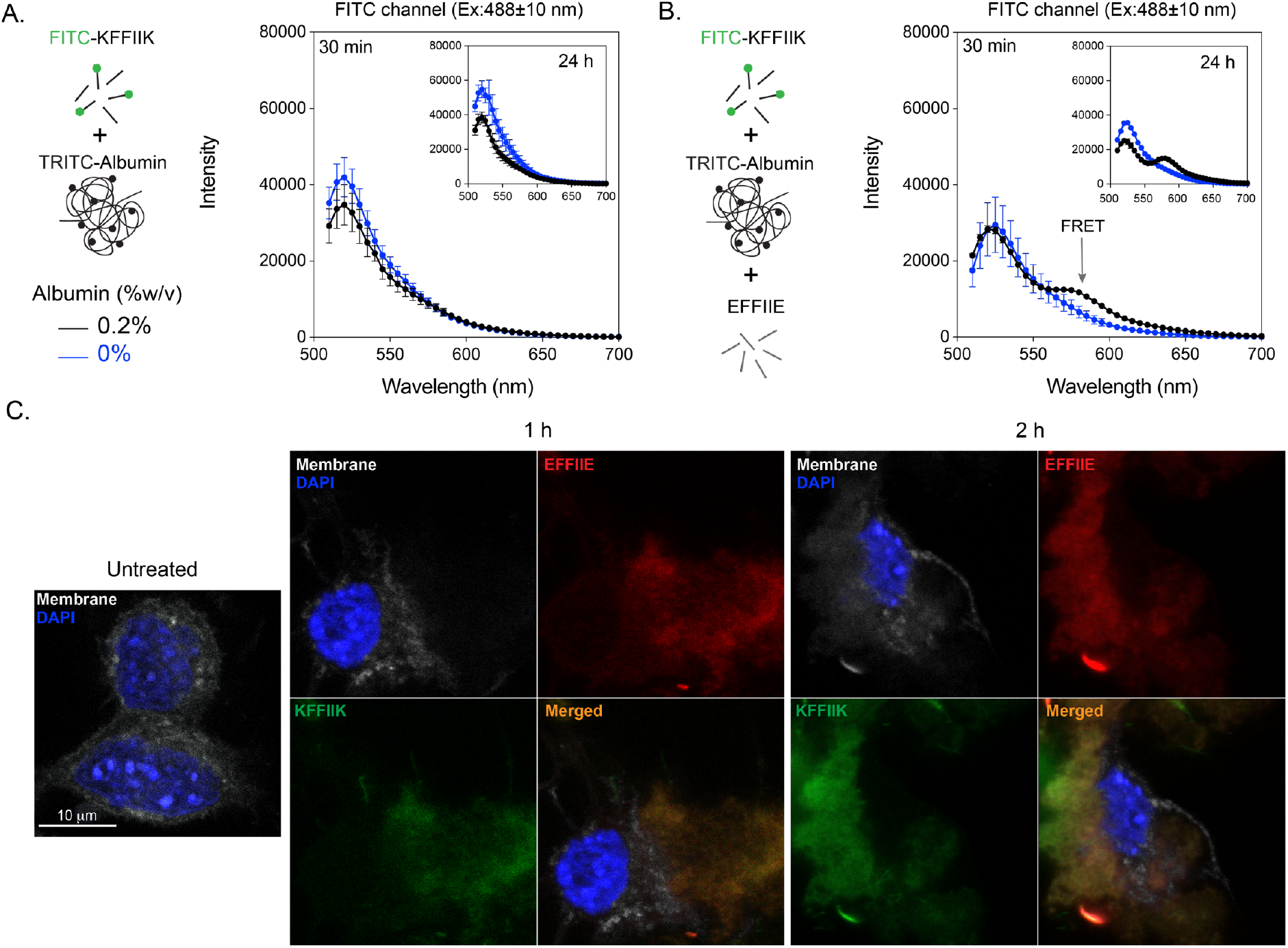
Albumin and [II] interaction. (A) The FRET analysis of the presence and absence of TRITC-albumin with FITC-KFFIIK and (B) with [II]. (C) Confocal analysis of cells treated with 0.5 mM [II] for 30 min, 1h, and 2h in the presence of 0.2 w% albumin. Membrane stained with WGA, gray; cells treated with Rhodamine B labeled EFFIIE, red; FITC labeled KFFIIK, green. (Scale bar = 10 μm)

The cell membranes were stained with Wheat Germ Agglutinin (WGA) (gray) and incubated with [II] (FITC-KFFIIK (green) and Rhodamine-EFFIIE (red)) in 0.2 w% albumin for 1h and 2h to identify the localization of the peptides. Accumulation of each peptide increased from 1h to 2h with lost integrity of the membrane, which also explained the higher LDH release **(Figure 3C)**. When the peptides were preincubated in the absence of albumin for 30 min, no cell membrane localization was observed **(Figure S8)**, also in line with viability results. Overall, these results demonstrates that in the absence of albumin, peptides assemble into ordered fibrillar structures and do not localize to the cell membrane. Presence of the albumin in the solution prevents immediate peptide-peptide interactions and allows time for peptide aggregation to occur within the cell and at the membrane. Therefore, changing albumin concentration is changing the peptide aggregation kinetics, which results in controllable membrane accumulation and cell death.

### 2.5 [II] aggregation induces cell stress

The cell plasma membrane senses external stimuli to adapt to changes through intracellular signaling. One of the pathways responding against a variety of external stimuli such as mechanical stress is the Hippo pathway. Aβ 1-42-mediated neurodegeneration involves the activation of the Hippo pathway.^31^ One of the central proteins within the Hippo pathway is YAP. YAP is a transcription factor found in the nucleus in active form and gets inactivated when translocate to the cytoplasm. We treated the cells to analyze the effect of [II] aggregation on YAP localization. At the early time points, 30 min, YAP localized in the cytoplasm **(Figure 5A)**. More specifically, cytoplasmic localization of YAP is mediated by its phosphorylation (Serine 127 (S127)), leading to its inactivation^32^ and degradation.^33^ Upon administration of [II] with 0.2 w% albumin, YAP phosphorylation in the cytoplasm can be observed green at 30 min and 1h **(Figure 5B)**. However, after 1h, phosphorylated YAP in the cytoplasm starts to degrade, which is parallel with YAP localization to the cytoplasm at early time points (30 min). Western blot results also confirmed the phosphorylation of YAP and degradation **(Figure 5C)**. Together these results show that [II] aggregation is inducing cytoplasmic localization of YAP through its phosphorylation, thus related to the mechanical stress on the cell membrane.

**Figure 5.**
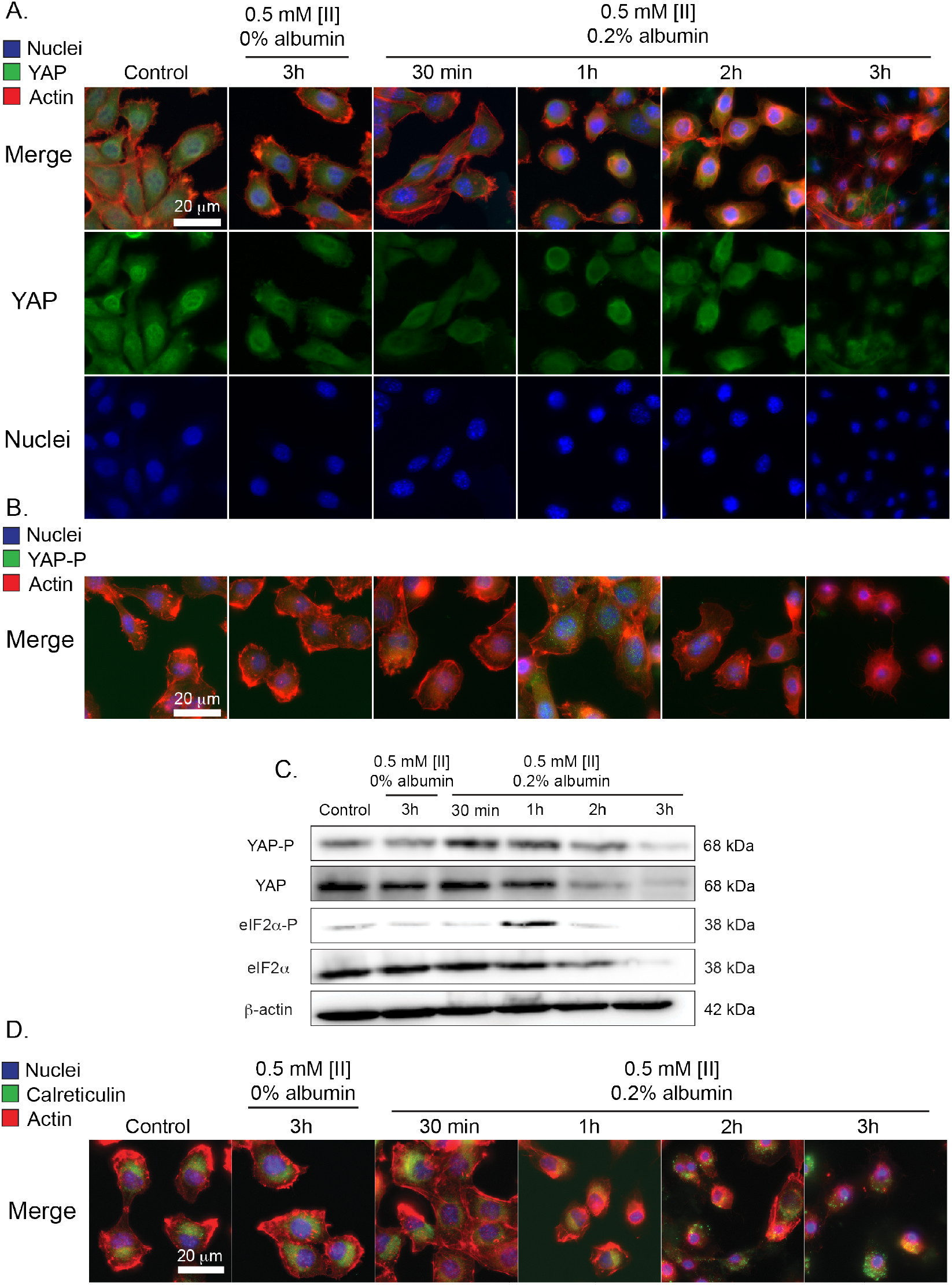
[II] aggregation on the cell membrane induces cell stress. (A) Immunocytochemistry of YAP localization (YAP = green, Actin = red, Nuclei = blue) (B) and phosphorylation based on aggregation (YAP-P = green). (C) Western blot analysis of YAP, YAP phosphorylation, eIF2α, and eIF2α phosphorylation in different time points, β-actin was used as an internal loading control. (D) Immunocytochemistry of calreticulin cell surface localization (calreticulin = green, Actin = red, Nuclei = blue). (Scale bars = 20 μm)

EIF2α is another factor integrated into the stress response to external stimuli. Under cell stress, eIF2α gets phosphorylated and initiates the activation of genes responsible for stress response. Phosphorylation of eIF2α is a consistent marker in a variety of neurodegenerative diseases,^34^ including AD.^35^ Histological analysis on AD patients showed elevated phosphorylation of eIF2α.^36^ Moreover, in in vitro studies, Aβ 1-42 was shown to induce eIF2α phosphorylation in human neuroblastoma cells.^37^ We analyzed eIF2α phosphorylation for 0.5mM [II] in 0.2w% albumin and 0.5mM [II] in 0% albumin (non-membrane damaging condition). [II] aggregation-induced eIF2α phosphorylation at 1h (Figure 5C) followed by rapid degradation due to proteasome activity reported previously.^38^

Phosphorylation of eIF2α and its downstream effect on calreticulin cell surface localization are identified as hallmarks of ICD.^39^ Cell membrane-localized calreticulin contributes to the immunogenicity of dying cancer cells, serving as an “eat me” signal to facilitate phagocytosis.^40,41^ We previously showed that [II] aggregation-induced cell death is ICD through the release of DAMPs in multiple cell lines.^12^ Given that eIF2α phosphorylation results in calreticulin surface localization, we identified the localization of calreticulin within the cells incubated with [II]. Calreticulin was observed only intracellularly in control cells and cells treated with 0.5mM [II] in 0% albumin. However, calreticulin localized to the cell membrane starting at 2h in cells treated with 0.5mM [II] in 0.2 w% albumin. These results show that [II] aggregation-induced cell death involves the phosphorylation of eIF2α, followed by calreticulin surface localization, indicates that the observed ICD is due to the mechanical stress on the cell membrane with [II] aggregation.

## 3. Conclusions

Here we show that [II] aggregation induces cell membrane damage and engages cell stress markers as in a variety of neurodegenerative disorders. Furthermore, we showed that this aggregation can be controlled by using albumin. [II] aggregation-induced the phosphorylation of eIF2α and calreticulin surface localization, which highlighted ICD. Overall, the modulation of peptide aggregation presented here, allows us to control cell membrane damage and offers us to modulate immunogenic cell death.

Protein and peptide aggregation is a phenomenon that can be seen in both pathogenic conditions and during developmental stages of amyloid proteins.^2,3^ Oligomeric species form transiently during the aggregation process and not only act as essential intermediates but also represent major pathogenic agents in disorders.^4^ However, the transient and complex structures and aggregates of oligomers create confusing results and challenge in drug development.^4,10^ In this study, we showed an engineered peptide complex via CoOP strategy as a synthetic and simple tool to study the amyloid aggregation mechanism into ordered assemblies (fibrils) and disordered aggregates (oligomers) with the presence of albumin, and correlate it to its toxicity. Importantly, we showed that, albumin can control the aggregation kinetics of [II], which induce mechanical stress on the cells and ICD. The immune response against ICD is potent. Controlling ICD means control over the immune response and progress of amyloidosis-related diseases.

Overall, [II] offers a unique engineered platform to study aggregation kinetics of amyloidosis. The simplicity of [II] as an engineered tool allowed the detailed characterization of disorganized structures to fibril aggregation kinetics and the effects of each step on cell. Studies with [II] can serve to establish clear and direct results for drug development with amyloidosis.

## 4. Experimental Section

### 4.1 Peptide Synthesis

KFFIIK and EFFIIE peptides were synthesized via solid phase peptide synthesis with PreludeX automatic peptide synthesizer (Protein Technologies, Inc., Tucson, AZ). They were prepared on a 0.25 scale by repeated amino acid couplings using Fmoc protected amino acid (3 eq. 1mL), DIC (7.5 eq. 1mL) and Oxyma (7.5 eq. 1mL). MBHA Rink Amide resin (High-Loaded, Gyros Protein Technologies) was used as solid support to construct the peptides. Fmoc protected amino acids (Gyros Protein Technologies) were removed through treatment with 20% piperidine/DMF solution for 45 min (three times for 15 min) at 80°C. All the peptide conjugation was performed for 2h at 80°C and acetylated with a treatment with 10% acetonitrile/DMF solution for 15 min twice.

Cleavage of the peptides from resin and deprotection of acid labile protected amino acids were carried out with a mixture of TFA/ TIS/ water in a ratio of 95:2.5:2.5 for 3 h at room temperature. Excess TFA and organic solvents were removed by evaporation and the remaining peptide was precipitated using diethyl ether at 80°C by using 1:4 volume ratio and stayed overnight. After precipitation, peptides in ether solutions were centrifuged at 8000 rpm for 15 min. Peptide precipitates were collected and ether was removed. The centrifuged white peptide precipitate was dissolved in either (0.1% formic acid in water for KFFIIK peptide) or (0.1% ammonium hydroxide in water for EFFIIE peptide). Peptides were purified with preparative HPLC (Agilent 1260) with Agilent ZORBAX 300 SB-C18 (9.4 x 250 mm) column with a mobile phase of water/acetonitrile mixture (0.1% ammonium hydroxide) used for negatively charged peptides; water/acetonitrile mixture (0.1% formic acid) was used for positively charged peptides. All peptides were tested with a purity >95%. HPLC run started with 100% water for 3 min, followed by a gradient increase in acetonitrile from 0% to 100% over 25 min, followed 100% acetonitrile for 3 min, and ended with 100% for 2 min. The flow rate is 2 mL/min and injection volumes are 10 μL.

### 4.2 Liposome formation and imaging

Giant liposomes were formed based on previously published data with simple modifications.^21^ First, agarose (UltraPure Agarose, ThermoFisher Scientific) was prepared with 1% w/v in water and boiled in a microwave oven until all agarose powder was dissolved. Then, it was poured down into a petri dish to cool down at room temperature to form a gel for 2-3h. Before coating the glass slides, agarose gel was melted in a microwave until all gel turned into a solution. One side of glass slides was dipped into agarose solution and excess solution was removed. After coating, glass slides were placed on a hotplate (37°C) and waited for 1-2h until an agarose film was formed on top of the glass slides. For making liposomes, DPPC (1,2-dipalmitoyl-sn-glycero-3-phosphocholine, Avanti Polar Lipids, Inc.) was used with cholesterol (Sigma-Aldrich). At first, DPPC was prepared in 2 mM in chloroform, and cholesterol was prepared in 400 μm in chloroform. Both of the solutions were mixed with equal volume (1 mL each) to have 5:1 molar ratio (DPPC: cholesterol). For staining the liposomes with DiR (1,1’-Dioctadecyl-3,3,3’,3’-Tetramethylindotricarbocyanine iodide) dye, 4 μL of DiR (1 mM in chloroform) was mixed with 2 mL DPPC: cholesterol. Then, 50 μL of labeled liposomes was dropped onto agarose-covered glass slides and spread out homogeneously by rolling a glass rod. Then, slides were placed in a vacuum chamber for 30 min. After removing from the vacuum chamber, slides were placed in a clean petri dish and 1xPBS was added from the side of the petri dish to allow hydration and swelling of liposomes for 3h. For peptide-membrane localization assay, FITC labeled KFFIIK and Rhodamine-B labeled EFFIIE peptides (Biomatik Corporation) were used. KFFIIK and EFFIIE were first solubilized in 10 mM as a stock solution in water, which was diluted to 0.5 mM with 1x PBS. The fluorescent portion consisted of 1:100 molar ratio. For example, FITC-KFFIIK was 50 μM in 0.5 mM KFFIIK peptide as a final concentration. After liposome formation, 0.5 mM of fluorescent labeled one peptide counterpart (200 μL) were incubated with liposomes for 60 min, then oppositely charged 0.5 mM of fluorescent labeled peptide (200 μL) was added and incubated for another 60 min. The imaging was performed with Keyence BZ-X800 with BZ-X filter Cy7 (DiR labeled liposome), BZ-X filter GFP (FITC-KFFIIK), and BZ-X filter mCherry (Rho-EFFIIE).

### 4.3 Aggregation analysis

Thioflavin T (ThT) (Sigma-Aldrich) and 1,6-Diphenyl-1,3,5-hexatriene (DPH) (Sigma-Aldrich) were used to understand the kinetics involved in ordered assembly and aggregation kinetics. Both fluorescence measurements included albumin (bovine serum albumin, Sigma-Aldrich) with different concentrations indicated based on %weight/volume (%w) and without albumin (in 1x PBS). At first, fresh albumin (40mg/mL, 4 %w) was dissolved in 1x PBS and diluted into 2, 1, 0.5, 0.2, 0.1, 0.05, 0.02, 0.002 w% in 1x PBS. Then, stock peptides (10 mM in water) were diluted (0.5 mM) in different albumin concentrations, individually. For ThT measurements, 5 μL from 400 μM ThT in PBS was added to 195 μL of 0.5 mM KFFIIK and EFFIIE (either in different albumin concentrations or in 1x PBS), individually. Then, each peptide in the same albumin concentration was mixed and read with BioTek Neo2SM microplate reader for 4 h with 10 min intervals (Ex: 440, Em:480 with gain:90). For DPH analysis, stock DPH (1 mM in water) was diluted to 80 μM working concentration in 1x PBS. Similar to ThT assay, 5 μL from 80 μM DPH in 1x PBS was added to 195 μL of 0.5 mM KFFIIK and EFFIIE (either in different albumin concentrations or in 1x PBS), individually. Then, each peptide in the same albumin concentration was mixed and read with BioTek Neo2SM microplate reader for 4h with 10 min intervals (Ex: 360, Em:450 with gain:100). Albumin with different concentrations (without any peptide) was also analyzed for both assays and subtracted from [II]+Albumin data. For [II] in 0 w% albumin, 1x PBS (with ThT or DPH) was used as a background for subtraction. For ThT analysis, relative intensity values were calculated within each group.

### 4.4 FRET analysis

FITC and TRITC could act as the donor and acceptor of a FRET pair, respectively. [28] FITC-KFFIIK peptide was prepared as 500 μM (1:50 FITC-KFFIIK:KFFIIK molar ratio) and albumin was prepared as 2% w/v(20 mg/mL, 300 μM) mixed with TRITC-albumin as 1:100 molar ratio. EFFIIE peptide was prepared as 500 μM. For a group consisting of FITC-KFFIIK and TRITC-albumin, 50 μL of FITC-KFFIIK was mixed 10 μL TRITC-albumin and 50 μL of PBS was added to have a final volume 110 μL. For 0 w% albumin, 50 μL of FITC-KFFIIK was mixed with 60 μL of PBS. [II] was prepared as follows. For 0.2w% albumin. 50 μL of FITC-KFFIIK was mixed with 50 μL of EFFIIE and 10 μL of TRITC-albumin was added to have a final volume 110 μL. For 0% albumin, 50 μL of FITC-KFFIIK was mixed with 50 μL of EFFIIE and 10 μL of PBS was added. These mixtures were incubated for 30 min and 24h at room temperature and subjected to fluorescence measurements with an excitation of 488 nm for FITC as donor and scan between the emission of 500 and 700 nm for TRITC as acceptor.

### 4.5 Cell Culture

EMT6 cells were cultured in Roswell Park Memorial Institute (RPMI) 1640 Medium, supplemented with 10% FBS (Hyclone SH30910.03) and 1% antibiotics; penicillin (100 U/mL), and streptomycin (100 μg/mL) (Thermo Fisher 15240062) according to the manufacturer’s instructions. Cells were cultured in a humidified incubator at 37°C supplied with 5% CO2. T75 flasks (TPP 90076) were used for culturing and cells were passaged upon 85% confluency by using trypsin (Sigma 59418C). Cell culture media was changed every 2 days.

### 4.6 Peptide preparation for in vitro experiments

[II] peptides were prepared in either 0.2 w% albumin or 0 w% albumin for in vitro conditions. First, albumin was pre- pared as a stock solution in 2% (20 mg in 1 mL the RPMI medium). Then, was diluted 1:10 to reach 0.2w% in the RPMI medium. The prepared KFFIIK and EFFIIE stock peptides (10 mM in water) were diluted to 0.5 mM separately into 0.2w% albumin/medium. For 0w% albumin condition, KFFIIK and EFFIIE stock peptides 10 mM in water) were diluted to 0.5 mM separately into albumin/medium. For 96 well plate, each well consisted of 100 μL of peptide having 50 μL EFFIIE and 50 μL KFFIIK in either only RPMI medium (0 w% albumin) or 0.2 w% albumin in RPMI medium.

### 4.7 Cell Titer Glo2.0 Viability

Cells were seeded onto 96 well plates and left for O/N attachment. The next day, media was removed and treatments were carried out. Viability was measured with Cell Titer Glo 2.0 solution (Promega G9248). The luminescent signal was measured in accordance with the manufacturer’s instructions. Measurements were carried out with BioTek Neo2SM microplate reader and relative viability was calculated by using untreated control group.

### 4.8 LDH Release

LDH release was measured with Cytoscan-LDH Cytotoxicity Assay (G-BIOSCIENCES 786-210). Briefly, at the end the treatments, collected supernatants were mixed with reaction mixture and incubated for 20 min at 37°C. Reaction was stopped with stop solution, and absorbance was measured at 490 nm and 680 nm. Triton-X treatment is used as a 100% LDH release control. Absorbance at 680 nm is used as the background signal and values were subtracted from absorbance values at 490 nm. Measurements were carried out with BioTek Neo2SM microplate reader and relative LDH release was quantified based on the positive control for LDH release (maximum LDH release through Triton-X treatment).

### 4.9 Confocal Imaging

EMT6 cells were seeded on glass coverslips in a 24 well plate and incubated overnight for attachment. Prior to the experiment, cells were labeled with WGA 633 membrane stain (ThermoFisher W21404) accordingly with the manufacturer’s instructions. After membrane staining, cells were washed 2x with PBS and treated in the presence (0.2 w%) and absence (0 w%) of albumin.

### 4.10 Immunoblotting, Reagents

Acrylamide/Bis-acrylamide, 30 w% solution (Sigma A3699), 1.5 M Tris-HCl, pH 8.8 (Teknova T1588), Tris HCl Buffer 0.5 M solution, sterile pH 6.8 (Bio Basic SD8122), Ammonium persulfate (Sigma A3678), UltraPure 10% SDS (Invitrogen 15553-027), TEMED (Thermo Fisher Scientific 17919), Dithiothreitol (DTT) (BIO-RAD 1610610) Tris Base (Fisher Bioreagents BP152), Glycine (Fisher Bioreagents BP381), 4x Laemmli sample buffer (BIO-RAD 1610747), TWEEN 20 (Sigma P9416), Mini Trans-Blot filter paper (BIO-RAD 1703932), Nitrocellulose Membranes 0.45 μm (BIO-RAD 1620115), EveryBlot Blocking Buffer (BIO-RAD 12010020), Clarity Western ECL Substrate (BIO-RAD 170-5060).

### 4.11 Procedure

Samples were prepared in Laemmli buffer and boiled for 8 min at 96°C. Then, proteins were separated with sodium dodecyl sulfate-polyacrylamide gel electrophoresis (SDS-PAGE) gels (8.5 w% and 15 w%) and transferred to nitrocellulose membranes (45 μm). After transfer, the membranes were blocked EverBlot Blocking Buffer and washed with Tris Buffered Saline with 0.1% Tween 20 (TBST). Blots were incubated with Phospho-YAP (Ser127) (Cell Signaling 13008), YAP (Cell Signaling 4912), Phospho-eIF2α (S51) (Cell Signaling 9721), eIF2α (Cell Signaling 9722) overnight. The next day blots were washed and incubated with Goat anti rabbit secondary antibody (Invitrogen 31460) and imaged with Clarity Western ECL substrate (BIO RAD 170-5060). β-actin (Santa Cruz sc-47778 HRP) was used as an internal loading control. Blots were imaged by using Azure c600.

### 4.12 Immunocytochemistry

EMT6 cells were seeded on glass coverslips in a 24 well plate and incubated overnight for attachment. After indicated treatments, samples were washed with 1X PBS, and fixed with 4% PFA for 20 min. Then, Triton-X (0.5%) was used to permeabilize the cells for 20 min on shaker. Samples were blocked with 3% BSA for 1h on shaker, washed 3 times with 1X PBS and incubated with the following antibodies Phospho-YAP (S127) (Cell Signaling 13008), YAP (Cell Signaling 4912) or calreticulin (Cell Signaling D3E6) overnight on shaker on ice. The next day samples were washed and incubated for 1h on shaker with DyLight™ 488 Donkey anti-rabbit IgG (minimal x-reactivity) Antibody (BioLegend 406404). After the incubation period, samples were washed and incubated with Flash Phalloidin™ Red 594 (BioLegend 424203) for 30 min on shaker. Lastly, samples were washed and mounted in ProLong™ Glass Antifade Mountant with NucBlue™ Stain (Thermo Fisher P36981) and stored in the dark until imaging.

## Acknowledgements

This work is supported in part by a grant from the Research Council of the University of Oklahoma Norman Campus and in part by the Oklahoma Tobacco Settlement Endowment Trust awarded to the University of Oklahoma, Stephenson Cancer Center. The content is solely the responsibility of the authors and does not necessarily represent the official views of the Oklahoma Tobacco Settlement Endowment Trust. Dr. Tingting Gu from Samuel Roberts Noble Microscopy Laboratory at the University of Oklahoma conducted the confocal imaging. EMT6 breast cancer cell line was a gift from Dr. Judy Lieberman, Boston Children’s Hospital, Harvard Medical School.

## Supporting Information

## References

1. Iadanza MG, Jackson MP, Hewitt EW, Ranson NA, Radford SE. A new era for understanding amyloid structures and disease. Nature Reviews Molecular Cell Biology. 2018;19(12):755–773. doi:10.1038/s41580-018-0060-8

2. Gallardo R, Ranson NA, Radford SE. Amyloid structures: much more than just a cross-β fold. Current Opinion in Structural Biology. 2020;60:7–16. doi:10.1016/j.sbi.2019.09.001

3. Jia Z, Schmit JD, Chen J. Amyloid assembly is dominated by misregistered kinetic traps on an unbiased energy landscape. Proceedings of the National Academy of Sciences. 2020;117(19):10322–10328. doi:10.1073/pnas.1911153117

4. Kulenkampff K, Wolf Perez AM, Sormanni P, Habchi J, Vendruscolo M. Quantifying misfolded protein oligomers as drug targets and biomarkers in Alzheimer and Parkinson diseases. Nat Rev Chem. 2021;5(4):277–294. doi:10.1038/s41570-021-00254-9

5. Fusco G, Chen SW, Williamson PTF, et al. Structural basis of membrane disruption and cellular toxicity by α-synuclein oligomers. Science. 2017;358(6369):1440–1443. doi:10.1126/science.aan6160

6. Yasumoto T, Takamura Y, Tsuji M, et al. High molecular weight amyloid β1-42 oligomers induce neurotoxicity via plasma membrane damage. Alzheimer’s & Dementia. 2020;16(S2):e037546. doi:https://doi.org/10.1002/alz.037546

7. Milner MT, Maddugoda M, Götz J, Burgener SS, Schroder K. The NLRP3 inflammasome triggers sterile neuroinflammation and Alzheimer’s disease. Current Opinion in Immunology. 2021;68:116–124. doi:10.1016/j.coi.2020.10.011

8. Tanaka H, Homma H, Fujita K, et al. YAP-dependent necrosis occurs in early stages of Alzheimer’s disease and regulates mouse model pathology. Nat Commun. 2020;11(1):507. doi:10.1038/s41467-020-14353-6

9. McKenzie BA, Dixit VM, Power C. Fiery Cell Death: Pyroptosis in the Central Nervous System. Trends in Neurosciences. 2020;43(1):55–73. doi:10.1016/j.tins.2019.11.005

10. Walsh DM, Selkoe DJ. Amyloid β-protein and beyond: the path forward in Alzheimer’s disease. Current Opinion in Neurobiology. 2020;61:116–124. doi:10.1016/j.conb.2020.02.003

11. Hamsici S, White AD, Acar H. Peptide framework for screening the effects of amino acids on assembly. Science Advances. Published online January 2022. doi:10.1126/sciadv.abj0305

12. Gunay G, Hamsici S, Lang GA, Lang ML, Kovats S, Acar H. Peptide Aggregation Induced Immunogenic Rupture (PAIIR). Advanced Science. 2022;n/a(n/a):2105868. doi:10.1002/advs.202105868

13. Moujalled D, Strasser A, Liddell JR. Molecular mechanisms of cell death in neurological diseases. Cell Death Differ. 2021;28(7):2029–2044. doi:10.1038/s41418-021-00814-y

14. Leong YQ, Ng KY, Chye SM, Ling APK, Koh RY. Mechanisms of action of amyloid-beta and its precursor protein in neuronal cell death. Metab Brain Dis. 2020;35(1): 11–30. doi:10.1007/s11011-019-00516-y

15. Finn TE, Nunez AC, Sunde M, Easterbrook-Smith SB. Serum Albumin Prevents Protein Aggregation and Amyloid Formation and Retains Chaperone-like Activity in the Presence of Physiological Ligands. Journal of Biological Chemistry. 2012;287(25):21530–21540. doi: 10.1074/jbc.M112.372961

16. Zhao M, Guo C. Multipronged Regulatory Functions of Serum Albumin in Early Stages of Amyloid-β Aggregation. ACS Chem Neurosci. 2021;12(13):2409–2420. doi:10.1021/acschemneuro.1c00150

17. Picón-Pagès P, Bonet J, García-García J, et al. Human Albumin Impairs Amyloid β-peptide Fibrillation Through its C-terminus: From docking Modeling to Protection Against Neurotoxicity in Alzheimer’s disease. Computational and Structural Biotechnology Journal. 2019;17:963–971. doi:10.1016/j.csbj.2019.06.017

18. Kakinen A, Javed I, Faridi A, Davis TP, Ke PC. Serum albumin impedes the amyloid aggregation and hemolysis of human islet amyloid polypeptide and alpha synuclein. Biochimica et Biophysica Acta (BBA) - Biomembranes. 2018;1860(9):1803–1809. doi:10.1016/j.bbamem.2018.01.015

19. Savini F, Bobone S, Roversi D, Mangoni ML, Stella L. From liposomes to cells: Filling the gap between physicochemical and microbiological studies of the activity and selectivity of host-defense peptides. Peptide Science. 2018;110(5):e24041. doi:10.1002/pep2.24041

20. Andrade S, Loureiro JA, Pereira MC. The Role of Amyloid β-Biomembrane Interactions in the Pathogenesis of Alzheimer’s Disease: Insights from Liposomes as Membrane Models. ChemPhysChem. 2021;22(15):1547–1565. doi:10.1002/cphc.202100124

21. Horger KS, Estes DJ, Capone R, Mayer M. Films of Agarose Enable Rapid Formation of Giant Liposomes in Solutions of Physiologic Ionic Strength. J Am Chem Soc. 2009;131(5):1810–1819. doi:10.1021/ja805625u

22. Chibowski E, Szcześ A. Zeta potential and surface charge of DPPC and DOPC liposomes in the presence of PLC enzyme. Adsorption. 2016;22(4-6):755–765. doi:10.1007/s10450-016-9767-z

23. Arosio P, J. Knowles TP, Linse S. On the lag phase in amyloid fibril formation. Physical Chemistry Chemical Physics. 2015;17(12):7606–7618. doi:10.1039/C4CP05563B

24. Stanyon HF, Viles JH. Human Serum Albumin Can Regulate Amyloid-β Peptide Fiber Growth in the Brain Interstitium: IMPLICATIONS FOR ALZHEIMER DISEASE *. Journal of Biological Chemistry. 2012;287(33):28163–28168. doi:10.1074/jbc.C112.360800

25. Biancalana M, Koide S. Molecular mechanism of Thioflavin-T binding to amyloid fibrils. Biochimica et Biophysica Acta (BBA) - Proteins and Proteomics. 2010;1804(7):1405–1412. doi:10.1016/j.bbapap.2010.04.001

26. Sang JC, Lee JE, Dear AJ, et al. Direct observation of prion protein oligomer formation reveals an aggregation mechanism with multiple conformationally distinct species. Chem Sci. 2019;10(17):4588–4597. doi:10.1039/C8SC05627G

27. Ziaunys M, Sakalauskas A, Sneideris T, Smirnovas V. Lysozyme Fibrils Alter the Mechanism of Insulin Amyloid Aggregation. International Journal of Molecular Sciences. 2021;22(4):1775. doi:10.3390/ijms22041775

28. Makwana PK, Jethva PN, Roy I. Coumarin 6 and 1,6-diphenyl-1,3,5-hexatriene (DPH) as fluorescent probes to monitor protein aggregation. Analyst. 2011;136(10):2161–2167. doi:10.1039/C0AN00829J

29. Kumar P, Nagarajan A, Uchil PD. Analysis of Cell Viability by the Lactate Dehydrogenase Assay. Cold Spring Harb Protoc. 2018;2018(6):pdb.prot095497. doi:10.1101/pdb.prot095497

30. Lichlyter DJ, Grant SA, Soykan O. Development of a novel FRET immunosensor technique. Biosensors and Bioelectronics. 2003;19(3):219–226. doi:10.1016/S0956-5663(03)00215-X

31. Irwin M, Tare M, Singh A, et al. A Positive Feedback Loop of Hippo-and c-Jun-Amino-Terminal Kinase Signaling Pathways Regulates Amyloid-Beta-Mediated Neurodegeneration. Front Cell Dev Biol. 2020;8:117. doi:10.3389/fcell.2020.00117

32. Hao Y, Chun A, Cheung K, Rashidi B, Yang X. Tumor Suppressor LATS1 Is a Negative Regulator of Oncogene YAP. Journal of Biological Chemistry. 2008;283(9):5496–5509. doi:10.1074/jbc.M709037200

33. Abylkassov R, Xie Y. Role of Yes-associated protein in cancer: An update. Oncology Letters. 2016;12(4):2277–2282. doi:10.3892/ol.2016.4955

34. Bond S, Lopez-Lloreda C, Gannon PJ, Akay-Espinoza C, Jordan-Sciutto KL. The Integrated Stress Response and Phosphorylated Eukaryotic Initiation Factor 2α in Neurodegeneration. Journal of Neuropathology & Experimental Neurology. 2020;79(2):123–143. doi:10.1093/jnen/nlz129

35. Ohno M. Roles of eIF2Î± kinases in the pathogenesis of Alzheimerâ€™/s disease. Front Mol Neurosci. 2014;7. doi:10.3389/fnmol.2014.00022

36. Chang RCC, Wong AKY, Ng HK, Hugon J. Phosphorylation of eukaryotic initiation factor-2α (eIF2α) is associated with neuronal degeneration in Alzheimer’s disease: NeuroReport. 2002; 13(18):2429–2432. doi:10.1097/00001756-200212200-00011

37. Lee DY, Lee KS, Lee HJ, et al. Activation of PERK Signaling Attenuates Aβ-Mediated ER Stress. Feany MB, ed. PLoS ONE. 2010;5(5):e10489. doi:10.1371/journal.pone.0010489

38. Yerlikaya A, Kimball SR, Stanley BA. Phosphorylation of eIF2α in response to 26S proteasome inhibition is mediated by the haem-regulated inhibitor (HRI) kinase. Biochemical Journal. 2008;412(3):579–588. doi:10.1042/BJ20080324

39. Bezu L, Sauvat A, Humeau J, et al. eIF2α phosphorylation is pathognomonic for immunogenic cell death. Cell Death Differ. 2018;25(8):1375–1393. doi:10.1038/s41418-017-0044-9

40. Obeid M, Panaretakis T, Joza N, et al. Calreticulin exposure is required for the immunogenicity of gamma-irradiation and UVC light-induced apoptosis. Cell Death Differ. 2007;14(10):1848–1850. doi:10.1038/sj.cdd.4402201

41. Fucikova J, Kepp O, Kasikova L, et al. Detection of immunogenic cell death and its relevance for cancer therapy. Cell Death Dis. 2020;11(11):1–13. doi:10.1038/s41419-020-03221-2

