## Supporting information for "Controllable membrane damage by tunable peptide aggregation with albumin"

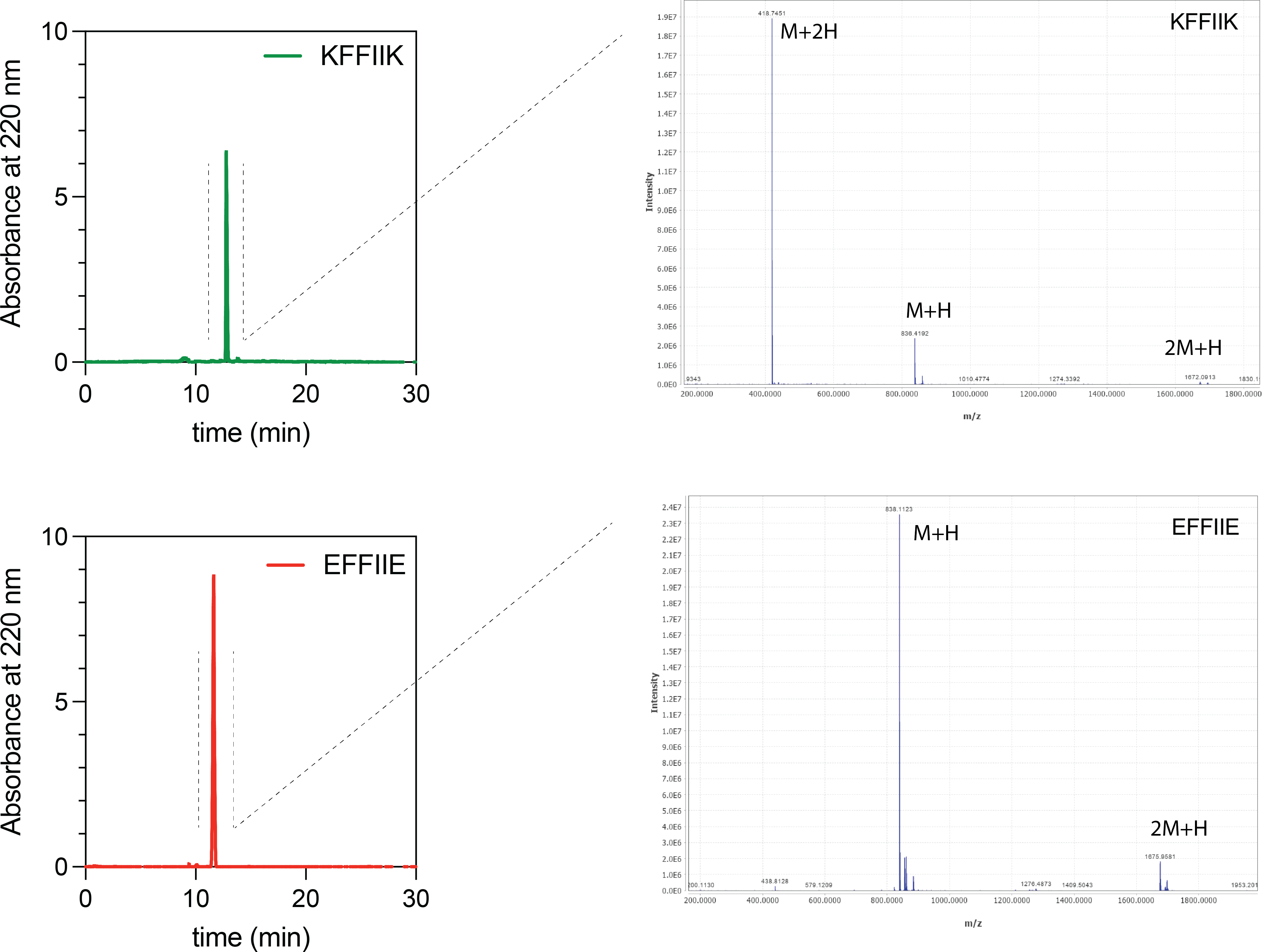

Figure S1. LC and MS analysis of KFFIIK and EFFIIE peptides

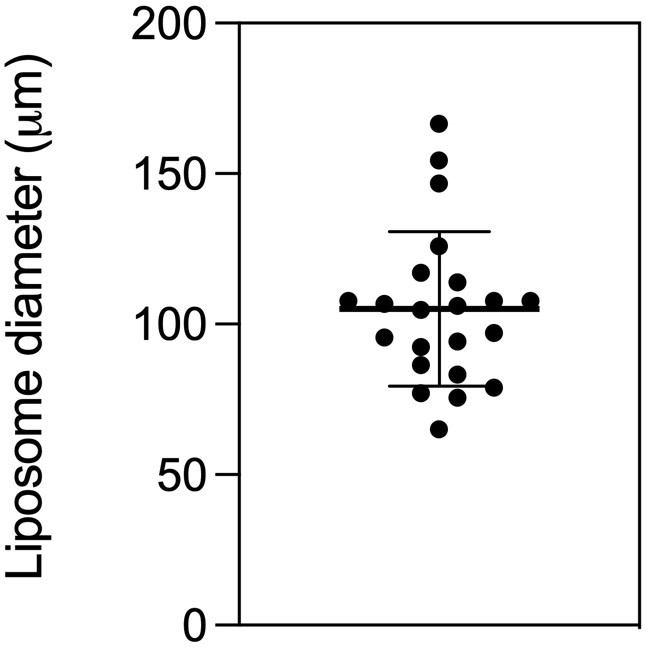

Figure S2. Films of agarose enable rapid formation of giant liposomes having around 100 µm in diameter

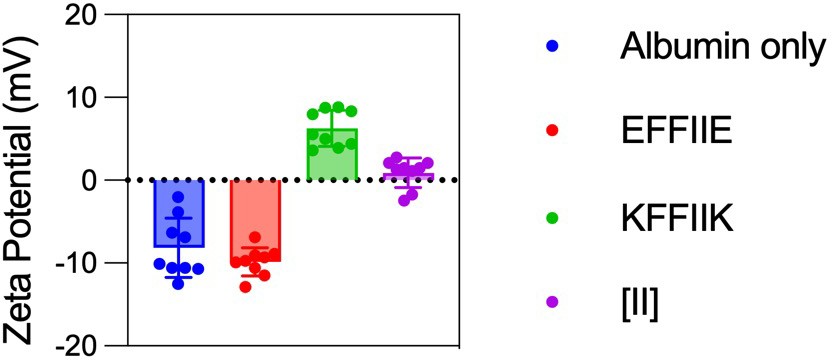

Figure S3. Zeta potential of individual oppositely charged peptides, co-assembled form and, albumin only

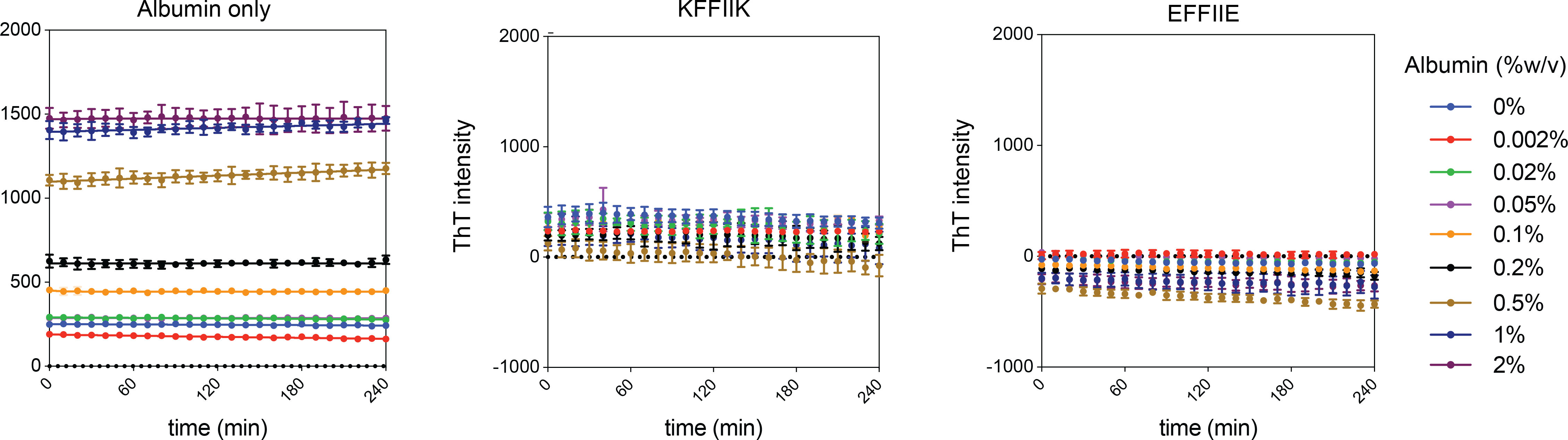

Figure S4. ThT analysis of albumin and individual peptides in diﬀerent concentrations

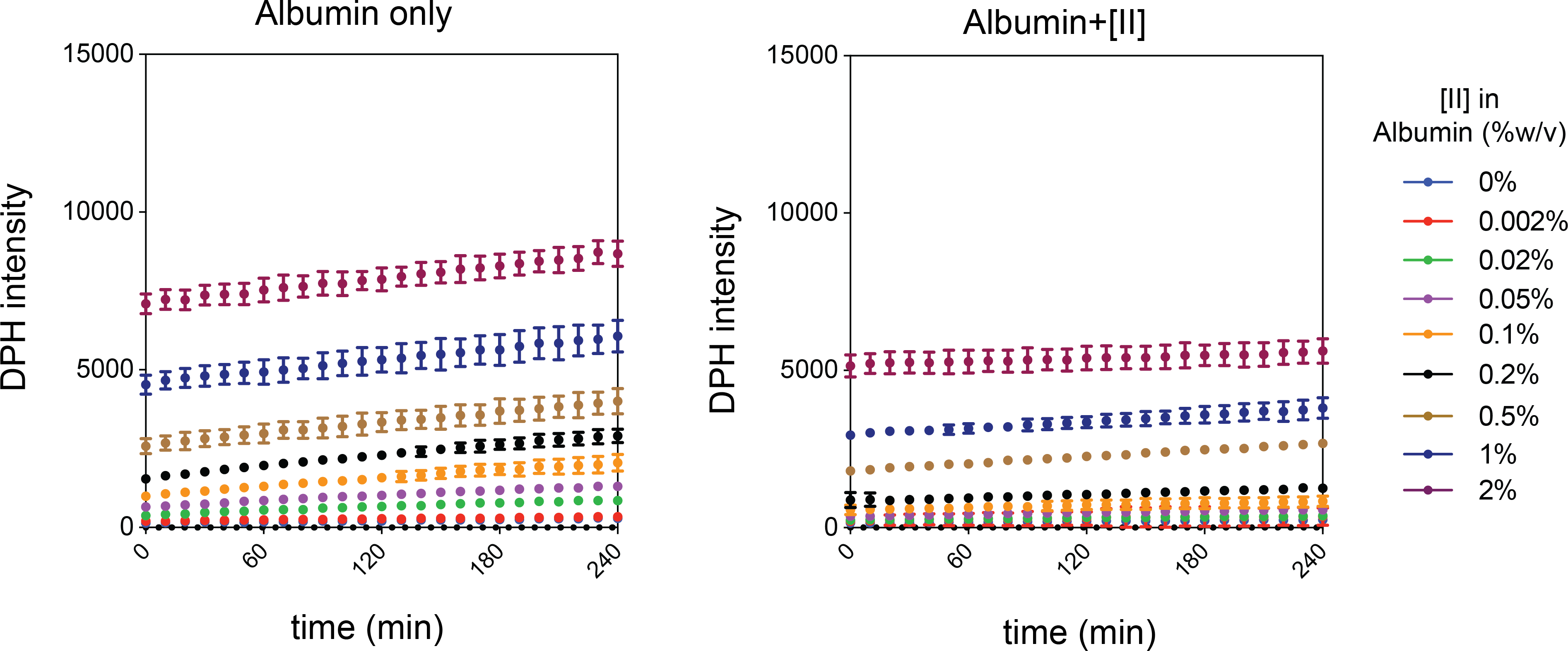

Figure S5. DPH analysis of albumin and albumin+[II] in diﬀerent concentrations

Figure S6. DPH analysis of individual peptides in diﬀerent albumin concentrations
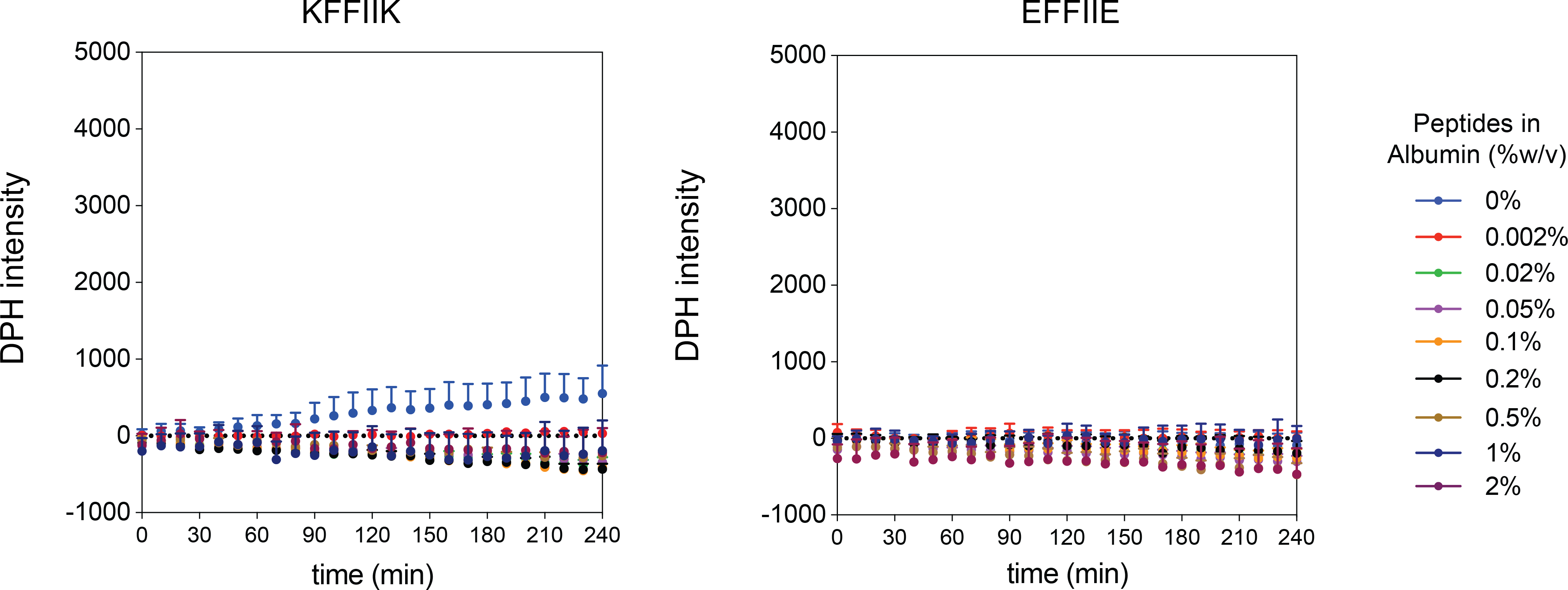

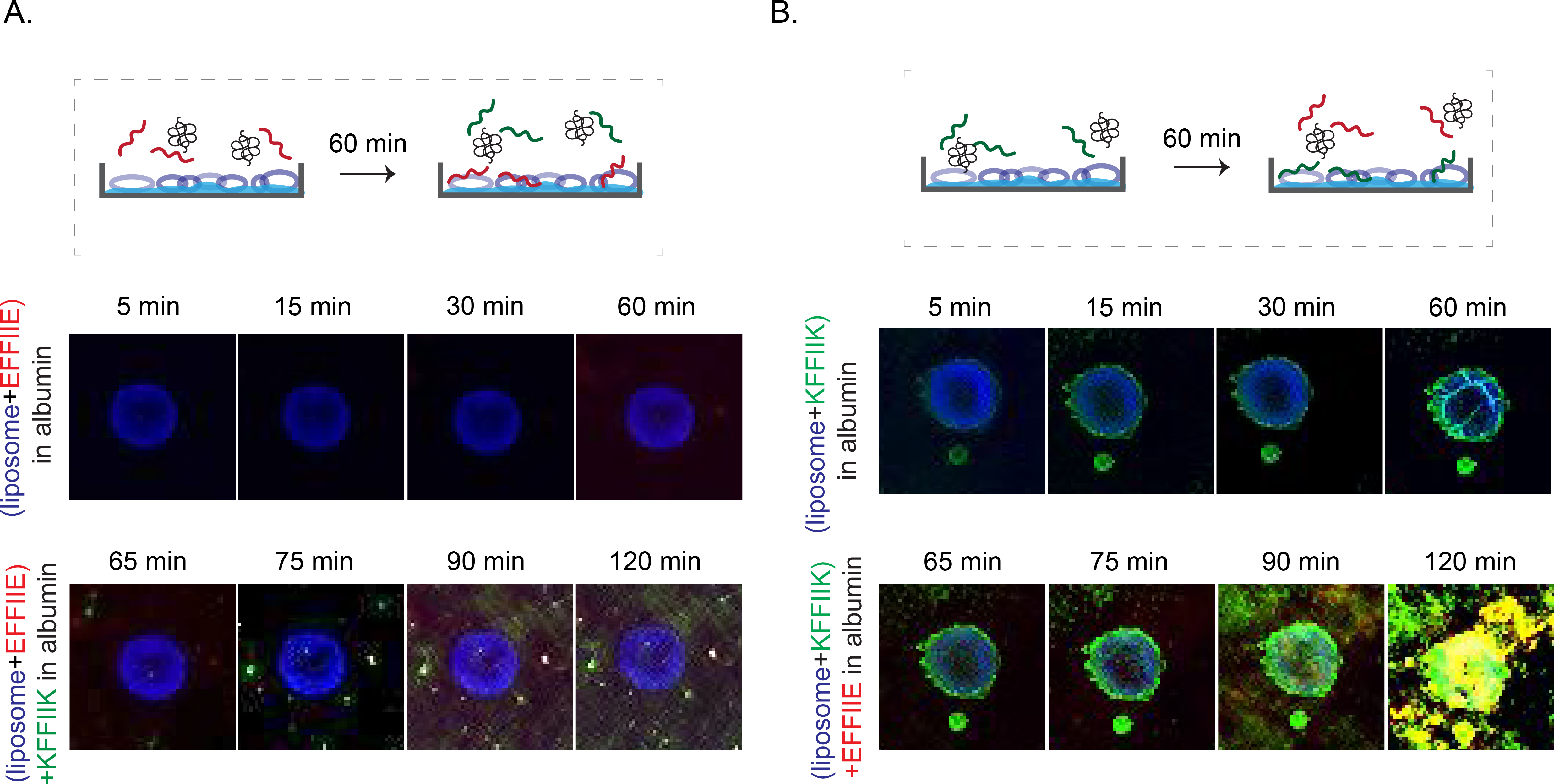

Figure S7. Membrane localization of individual peptides with the presence of albumin

Figure S8. The treatment of cells with preincubated FITC and Rhodamine B labelled [II] without albumin for 1 h and 2 h
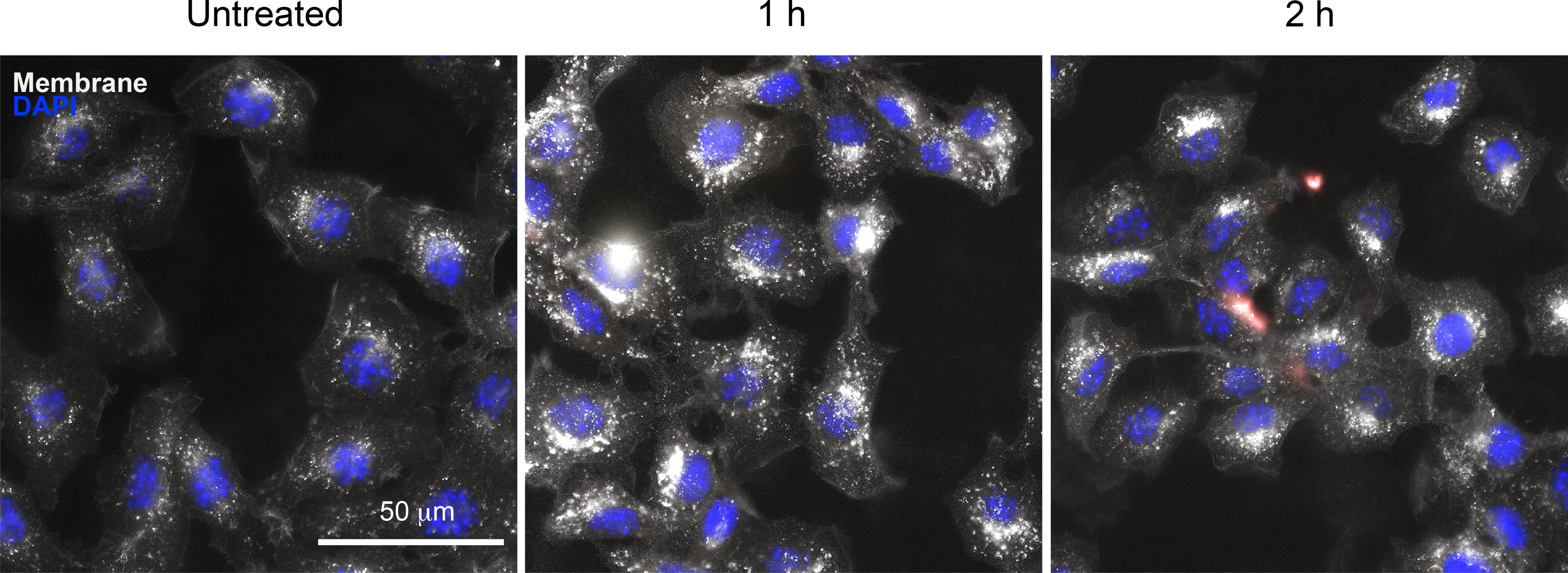
. Membrane was stained with WGA, white.
